# Addiction associated N40D mu-opioid receptor variant modulates synaptic function in human neurons

**DOI:** 10.1101/328898

**Authors:** Apoorva Halikere, Dina Popova, Aula Hamod, Mavis R. Swerdel, Jennifer C. Moore, Jay A. Tischfield, Ronald P. Hart, Zhiping P. Pang

## Abstract

**Background:** The OPRM1 A118G gene variant (N40D) encoding the µ-opioid receptor (MOR) has been associated with dependence on opiates and other abused drugs but its mechanism is unknown. With opioid abuse-related deaths rising at unprecedented rates, understanding these mechanisms may provide a path to therapy.

**Methods:** Seven human induced pluripotent stem (iPS) cell lines from homozygous N40D subjects (4 with N40 and 3 with D40 variants) were generated and human induced neuronal cells (iNs) were derived from these iPS cell lines. Morphological, gene expression as well as synaptic physiology analyses were conducted in human iN cells carrying N40D MOR variants; Two pairs of isogenic pluripotent stem cells carrying N40D were generated using CRISPR/Cas9 genome-editing and iN cells derived from them were analyzed.

**Results:** Inhibitory human neurons generated from subjects carrying N40D MOR gene variants show mature properties in morphological and functional analyses. Gene expression revealed that they express mature neuronal marker and MORs. Activation of MORs suppressed inhibitory synaptic transmission in human neurons carrying both N40 or D40 MOR variants but D40 show stronger effects. To mitigate the confounding effects of background genetic variation on neuronal function, the regulatory effects of MORs on synaptic transmission were validated in two sets of independently generated isogenic N40D iNs.

**Conclusions:** Activations of N40D MOR variants show different regulatory effects on synaptic transmission in inhibitory human neurons. This study identifies neurophysiological differences between human MOR variants that may predict altered opioid responsivity and/or dependence in this subset of individuals.

## INTRODUCTION

Well over 46,000 Americans died of opioid overdose in 2016, with the sharp increase in 2014 – 2016 due to synthetic opioids (1), prompting a public health crisis whose biological underpinnings are poorly understood. The µ-opioid receptor (MOR) mediates the most powerful addictive properties of abused opiate alkaloids and much research has identified chemically diverse ligands of varying efficacies for pain relief or treatment of addiction. Because of its substantive role in mediating reward and positive reinforcement, MOR is also an indirect target of alcohol, nicotine, and other drugs of abuse (2, 3). MOR-mediated synaptic alterations in reward-associated brain regions may represent a key underlying mechanism of reinforcement in drug abuse (4), but our understanding of this process in human neurons is limited.

Human genetic studies suggest that MOR gene variants play key roles in susceptibility to opioid addiction in humans. Most prominently, the A118G single nucleotide polymorphism (SNP) in OPRM1, rs1799971, a non-synonymous gene variant which replaces asparagine at position 40 (N40) with aspartate (D40), is found in up to 50% of individuals in certain ethnic groups and is associated with drug dependence phenotypes (5). There have been a number of investigations (5–13) into the functional consequences of the MOR D40 variant on receptor activation in overexpression models, knock-in mice, and primate models, but no systematic investigations into the functional and electrophysiological consequences of OPRM1 A118G have been reported, specifically not in a human neuronal context. Understanding how the D40 variant affects MOR signaling and synaptic function when expressed at normal levels in human neurons may provide insight into mechanisms underlying drug abuse, at least in people carrying this variant.

In order to fill the gap in studies done in the mouse and heterologous systems, we generated human induced neuronal (iN) cells from induced pluripotent stem (iPS) cells derived from subjects carrying homozygous alleles for either MOR N40 or D40 in order to better dissect the role of MOR N40D in a physiologically relevant and human-specific model system. Strikingly, we found that D40 MOR human neurons exhibit a stronger suppression of inhibitory synaptic release in the presence of MOR-specific agonist DAMGO ([D-Ala^2^, N-MePhe^4^, Gly-ol]-enkephalin) compared to N40 human neurons. In order to control for the possibility of individual genetic background variation between subject cell lines, we used CRISPR/Cas9 gene targeting to generate two sets of isogenic human stem cell lines: one pair with a 118GG knock-in into a well-characterized human embryonic stem (ES) cell line and the other by converting a minor allele carrier (118GG, D40) into a major allele carrier (118AA, N40). Remarkably, the synaptic regulations of MOR activation in the isogenic lines recapitulate those of neurons generated from different human subjects. This study exemplifies the use of patient-specific iPS cells as well as gene targeted isogenic stem cell lines to advance our understanding of the fundamental cellular and synaptic alterations associated with MOR N40D in human neuronal context.

## METHODS AND MATERIALS

### Generation of human iPS cells from lymphocytes of subjects carrying MOR N40D

Human iPS cell lines were generated by RUCDR Infinite Biologics ^®^ from human primary lymphocytes carrying either MOR N40 or D40 genotypes using Sendai viral vectors (CytoTune^TM^, ThermoFisher Scientific), as previously described (14). Human iPS cells were cultured and maintained as described previously (15).

### Human iPS cell maintenance

Human iPS cells were cultured in 37°C, 5% CO_2_ on Matrigel^®^ Matrix (Corning Life Sciences)-coated plates in mTeSR medium (Stem Cell Technologies). For passaging and differentiation done weekly, iPS cells were dissociated using Accutase (Stem Cell Technologies), spun down at 1000 RPM for 5 minutes, and replated at a density of 20,000 cells/cm^2^ for maintenance cultures and 50,000 cells/cm^2^ for differentiation.

### Lentivirus preparation

Lentiviruses were produced in HEK293T cells by co-transfection of the three envelope proteins REV, RRE and VSVG vectors with 22µg of either FUW-Tet-O-Ascl1-T2A-puromycin, FUW-Tet-O-Dlx2-IRES-hygromycin, or FUW-rtTA. For each transfection, 9.1µg of REV, 13.77 µg VsVg, 19.1 µg RRE with 22g of lentiviral vector was transfected into a 150mm dish of HEK293T cells of 60% confluency using calcium phosphate transfection technique. Media was changed 12 hours following transfection, and virus was harvested in the media 48 hours following transfection, pelleted using an ultra-centrifuge (25,000 RPM for 2 hours), resuspended in MEM and aliquoted. Virus was stored in −80°C until use.

### Generations of isogenic human stem cell lines carrying N40D MOR gene variants

Two pairs of isogenic N40D MOR human stem cells lines were generated using CRISPR/Cas9 genome editing. Briefly, to convert H1 embryonic stem (ES) cells carrying homozygous AA118 major allele to GG118 homozygous minor alleles, a sgRNA designed from Optimized CRISPR Design Tool (http://crispr.mit.edu/) and Cas9 were expressed using the PX459 vector (Addgene plasmid #62988) and was transfected using Lipofectamine 3000 reagent (ThermoFisher Scientific, L300015) along with a single stranded oligodeoxynucleotide (ssODN) of 140 base pairs with homology arms flanking the mutation site carrying mutations for G118, a BamHI restriction enzyme site for screening, along with a mutation to mutate the PAM sequence. Individual clones were hand-picked for expansion and screening by PCR and sequencing. Heterozygous clone 9-2 was expanded and transfected for targeting the second allele of OPRM1 Exon 1. The two homozygous G118 knock-in clones were further subcloned before expansion and freezing.

To convert rs1799971 in the 03SF subject iPS cell line from homozygous minor allele (GG) to major allele (AA), a slightly different strategy was used. First, a CRISPR targeting site was found using ZiFit software (16). The target site (GGCAACCTGTCCGACCCATG) included the major allele sequence so the gRNA was designed to incorporate the minor allele (GGCgACCTGTCCGACCCATG). A 200 nt homologous recombination donor oligo was designed to convert minor to major allele, inactivate the CRISPR site, and introduce a HpaI site for screening. The gRNA was synthesized by PCR and in vitro transcription (GeneArt Precision gRNA Synthesis Kit, Life Technologies) (17), mixed with synthetic Cas9 protein (Life Technologies), donor oligo, and the mixture was electroporated into iPS cells (Amaxa nucleofector, Lonza) along with a GFP expression plasmid (pGFP-Max, Lonza). One day later, cells were dissociated with Accutase and GFP-expressing cells were collected by FACS and plated at about 5,000 cells per well in a 6-well plate on irradiated MEFs. By 7-10 days, colonies were visible and hand-picked for screening. Three iPS cell clones were selected: C12, which had no evidence of editing to be used as a negative control; D11 and A10, which both had homozygous edits to produce rs1799971 major allele (AA). In all gene-targeted cell lines, sequencing confirmed these edits and that all predicted off-target sites were unchanged.

### Generation of GABAergic iN cells from human ES and iPS cells

The protocol of generating GABAergic human iN cells was described recently (18). Briefly, iPS cells and ES cells were plated as dissociated cells on Matrigel ^®^ Matrix (Corning Life Sciences)-coated dishes in mTeSR (Stem Cell Technologies) medium with 2µM Y-27632 (Stemgent). The following day, the cells are infected with Ascl1, Dlx2 and rtTA lentiviruses for 10-12 hours upon which culture medium was replaced with Neurobasal medium (GIBCO by Life Technologies) with B27 and L-Glutamine supplemented with 2µg/mL Doxycycline (MP Biomedicals) and 2µM Y compound to induce TetO expression. The protocol for generating lentiviruses expressing different transcription factors was previously described (18). Puromycin and Hygromycin selection was conducted for the following 2 days, and on day 5, the iN cells are dissociated with Accutase and plated on glass coverslips with a monolayer of passage three primary astrocytes isolated from p1-3 pups, as described previously (15, 18, 19). Following plating, 50% of the culture medium was replaced every 2-3 days with fresh Neurobasal media containing B27, L-Glutamine, 100 ng/ml of BDNF, NT3 and GDNF.

### Real-time RT-PCR (qPCR)

Total neuronal RNA from three independently generated batches of iN cells for each cell line was prepared using TRIzol ^®^ Reagent (Thermo Fisher Scientific), Human-specific Taqman probes were purchased for OPRM1, MAP2, Tuj1, VGAT, GAD1, TH and PCR reaction conditions followed the manufacturer’s recommendations. Undifferentiated iPS cells, ES cells, and mouse astrocytes were used as negative controls. A sample of total RNA of a healthy human brain as well as Human Thalamus from Biochain ^®^ was used as a positive control. Relative RQ values were obtained by normalizing expression levels to the C12 iN condition. Student’s t-test was used to compare grouped N40 and D40 means.

### Immunocytochemistry and confocal imaging

Inhibitory human neurons were fixed for 15 minutes in 4% paraformaldehyde in PBS and permeabilized using 0.1% Triton X-100 in PBS for 10 minutes at room temperature. Cells were then incubated in blocking buffer (4% bovine serum albumin with 1% normal goat serum in PBS) for 1 hour at room temperature and then incubated with primary antibodies diluted in blocking buffer for 1 hour at room temperature, washed with PBS three times, and subsequently incubated in secondary antibodies for 1 hour at room temperature. Confocal imaging analysis was performed using a Zeiss LSM700. Primary Antibodies used include: mouse anti Oct4 (Millipore Sigma MAB4401, 1:2000), mouse anti Tra-1-60 (Millipore Sigma MAB4360, 1:1000), mouse anti MAP2 (Sigma-Aldrich M1406, 1:500), rabbit anti MAP2, (Sigma-Aldrich M3696, 1:500), rabbit anti Synapsin (e028, 1:3000), rabbit anti VGAT (Millipore Sigma AB5062P, 1:2000), mouse anti Gad-67 (Abcam ab26116, 1:500), mouse anti β3 Tubulin (BioLegend 801201, 1:2000).

### Electrophysiology

Functional analyses of iN cells were conducted using whole cell patch clamp as described elsewhere (15, 20). Briefly, a K-Gluconate internal solution was used, which consisted of (in mM): 126 K-Gluconate, 4 KCl, 10 HEPES, 4 ATP-Mg, 0.3 GTP-Na_2_, 10 Phosphocreatine. The pH was adjusted to 7.2 and osmolarity was adjusted to 270-290 mOsm. The bath solution consisted of (in mM): 140 NaCl, 5 KCl, 2 CaCl_2_, 2 MgCl_2_, 10 HEPES, 10 Glucose. The pH was adjusted to 7.4. Spontaneous inhibitory postsynaptic currents (sIPSCs) were recorded at a holding potential of 0mV under voltage-clamp mode. Miniature IPSCs were recorded in the presence of tetrodotoxin (1 µM). Intrinsic action potential firing properties of the iN cells were recorded in a bath solution containing 50 µM Picrotoxin and 20 µM CNQX. Evoked synaptic currents were elicited using an extracellular concentric bipolar stimulating electrode positioned approximately 100 µm away from the cell soma. All recordings were performed at room temperature. Data presentation: All data are presented as mean ± S.E.M. Student’s t-test or 2-way ANOVA were used to assess statistical significance.

## RESULTS

### Generation of human inhibitory neurons carrying N40D MOR variants

To investigate the functional role of the MOR N40D variant in a human neuronal context, we obtained iPS cells from multiple individuals of European descent carrying homozygous alleles for either MOR N40 (n=4) or MOR D40 (n=3) (**Supplemental Fig. 1A**). The rs1799971 genotype and the pluripotency of all seven iPS cell lines are confirmed by sequencing and colocalized immunocytochemistry (ICC) for OCT4 and Tra-1-60 (**Fig. 1A-B**).

**Figure 1.**
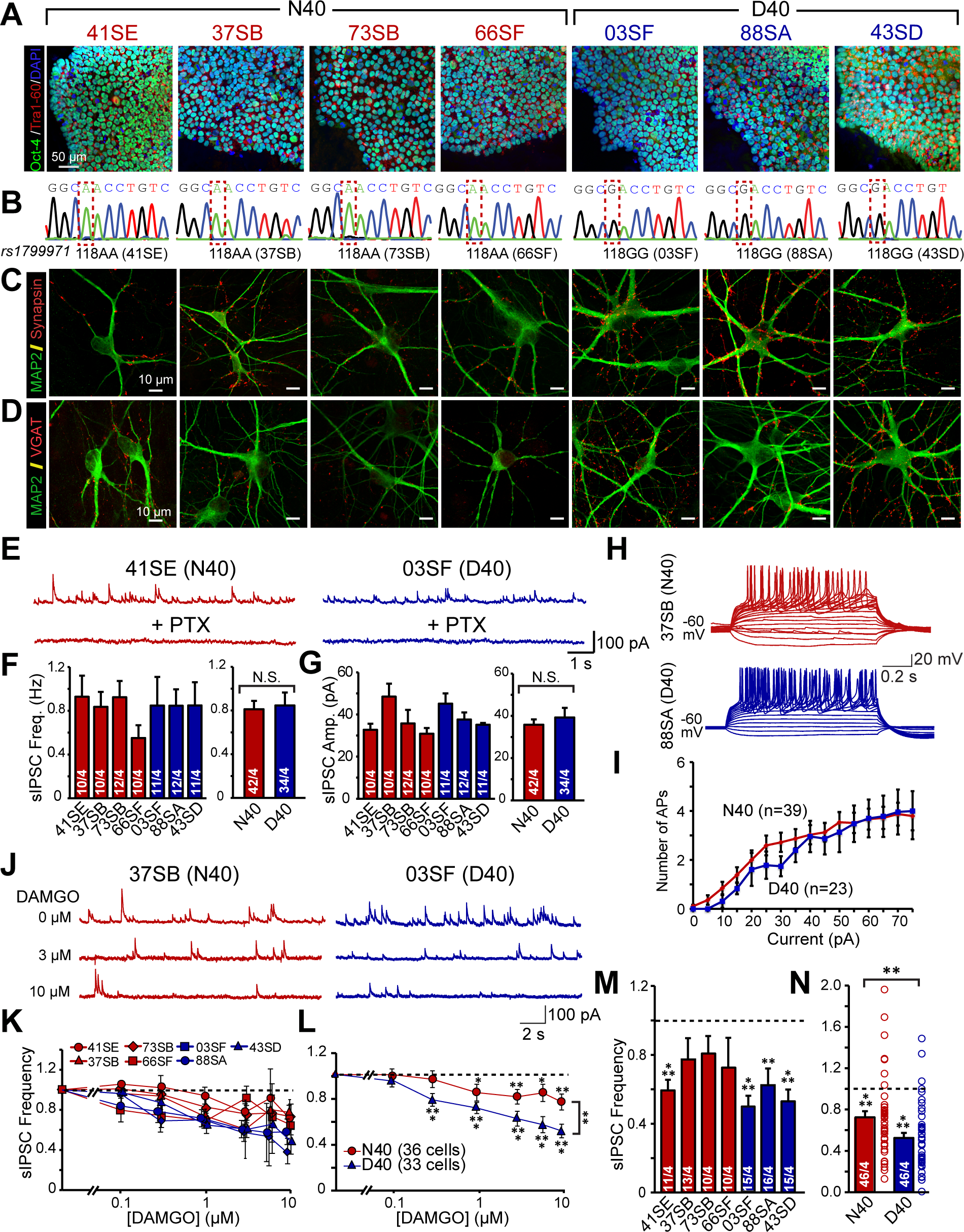
MOR N40D expressing inhibitory human neurons exhibit more robust suppression of inhibitory synaptic transmission by DAMGO. **(A)** Oct4 (green), Tra 1-60 (red), and Dapi (blue) ICC for N40 and D40 subject iPS cells depicting pluripotency **(B)** Sequencing confirming homozygous A118 or G118 genotype of human iPS cell lines **(C)** MAP2 (green) and Synapsin (red) ICC of iN cells generated from N40 and D40 subject iPS cells **(D)** MAP2 (green) and VGAT (red) ICC of induced inhibitory neuronal (iN) cells generated from N40 and D40 subject iPS cells **(E-G)** Both N40 and D40 iN cells exhibit PTX sensitive spontaneous IPSCs whose frequency (N40 vs D40: N.S.) and amplitude (N40 vs D40: N.S.) are unaffected by MOR N40D substitution **(H)** Representative traces of action potentials induced by step current injections (from −20 to +75 pA, 5pA increments) during current clamp recordings from one N40 and D40 cell line **(I)** Quantification of induced action potentials in inhibitory iNs cells illustrating that neuronal excitability is unchanged as a consequence of MOR N40D (N40 vs D40: N.S. at all current injections) **(J)** Representative traces of sIPSCs recorded to increasing concentrations of DAMGO in N40 and D40 iN cells **(K)** Quantification of inhibition of sIPSC frequency in individual subject derived N40 and D40 iN cells **(L)** Merged data of the four N40 and three D40 subject lines illustrates that D40 iN cells exhibit stronger suppression of IPSC frequency compared to N40 iN cells **(M-N)** sIPSC frequency response to a single concentration of 10µM DAMGO; data is normalized to control (N40 vs control: p<0.001, D40 vs control: p<0.001). Summary graphs are shown as individual cell lines or merged data of either four N40 patients (red bars) and three D40 patients (blue bars) (N40 vs D40: p <0.01). Data are depicted as means ± SEM. Numbers of cells/Number of independently generated cultures analyzed are depicted in bars. Statistical significance between N40 and D40 was evaluated by Student’s T test (*p<0.05, **p<0.01, ***p<0.001).

We derived inhibitory induced neuronal (iN) cells from all 7 iPS cell lines by lentiviral mediated ectopic expression of the transcription factors Ascl1 and Dlx2^33^. These induced human neuronal cells express pan-neuronal makers including MAP2, β3-tubulin, and Synapsin (**Fig. 1C, Supplemental Fig. 1B**) as well as inhibitory neuronal markers GAD67 and VGAT (**Fig. 1D, Supplemental Figs. 1C**). Thus, the N40D SNP has no impact on MOR expression or inhibitory neuronal identity. To examine whether the N40 and D40 iN cells are functionally comparable under baseline conditions, we performed whole cell patch-clamp recordings of iN cells after 5-6 weeks of re-plating onto a monolayer of mouse astroglia. The iN cells of both genotypes exhibit similar intrinsic membrane excitability (**Supplemental Fig. 1D-F**) and can fire repetitive spontaneous action potentials (APs) at baseline levels (**Supplemental Fig. 1G-I**) and exhibit similar intrinsic excitability under baseline conditions (**Fig. 1H-I**). Similarly, no significant differences in spontaneous or miniature inhibitory postsynaptic currents (sIPSCs and mIPSCs, respectively) were observed by genotype (**Figs. 1E-G, Supplemental Fig. 1J-L**), indicating that the N40D variant does not affect passive or active membrane properties and that the neurons generated from subject iPS cell lines are of similar functional maturation and differentiation.

### MOR D40 iN cells exhibit altered sensitivity to the MOR agonist DAMGO

There have been numerous studies^7, 22-24, 28, 36-39^ examining the functional consequences of MOR N40D on receptor activation in overexpression models and in knock-in mice harboring MOR N40D, but no functional or electrophysiological analyses on cultured neurons have been conducted, specifically not in a human neuronal context. To gauge whether N40 and D40 iN cells may respond differently to MOR activation, we used a MOR-specific agonist DAMGO ([D-Ala^2^, N-MePhe^4^, Gly-ol]-enkephalin) to study its role modulating synaptic release. In both N40 and D40 iN cells, DAMGO suppressed sIPSCs in a dose-dependent manner (**Fig. 1K**). However, the suppression of sIPSC frequency was more robust in D40 iN cells compared to N40 iN cells in multiple repeated experiments and multiple iPS cell lines (**Fig. 1L**), with no difference in sIPSC amplitude by genotype. To confirm that the observation is not due to a residual effect of prolonged agonist exposure, we applied a single concentration of 10µM DAMGO (**Fig. 1M-N**) and similarly found that D40 iN cells respond more robustly to MOR activation compared to N40 iN cells, illustrating genotype-dependent regulation of MOR signaling.

### Generation of isogenic human pluripotent stem cell lines carrying MOR N40D SNP

To directly compare the two MOR genotypes in identical genetic backgrounds, eliminating the impact of secondary genetic variation, we generated two sets of isogenic human stem cell lines using CRISPR/Cas9 gene targeting. We first targeted the MOR locus in a well-characterized human H1 ES cell line, which carries only major allele (A118), using an sgRNA targeting the antisense DNA strand along with a 140bp single stranded oligodeoxynucleotide carrying G118 (**Fig. 2A-B**). Simultaneously, we converted one iPS cell line with the homozygous G118 allele to a homozygous A118 genotype with an alternative strategy utilizing direct transduction of guide RNA and Cas9 protein into the subject cell line (**Fig. 2C-D**). We isolated two clones (**Supplemental Fig. 2A-C**) with no detectable off-target effects from each targeting scheme. Inhibitory iN cells generated from isogenic lines stain positive for MAP2, Synapsin and VGAT (**Fig. 2E-F**) and exhibit similar intrinsic membrane properties (**Supplemental Fig. 2D-E**) as well as sIPSC and AP properties in both genotypes (**Supplemental Figs. 2F-G**, **3A-C**). The densities and sizes of synapses were also not significantly different between genotypes (**Supplemental Fig. 2H-I**), illustrating that the N40D SNP does not alter synaptogenesis or functional maturation in the isogenic human neurons, and that the MOR N40D SNP has no consequence on iN cell maturation or synaptic transmission at baseline levels.

**Figure 2.**
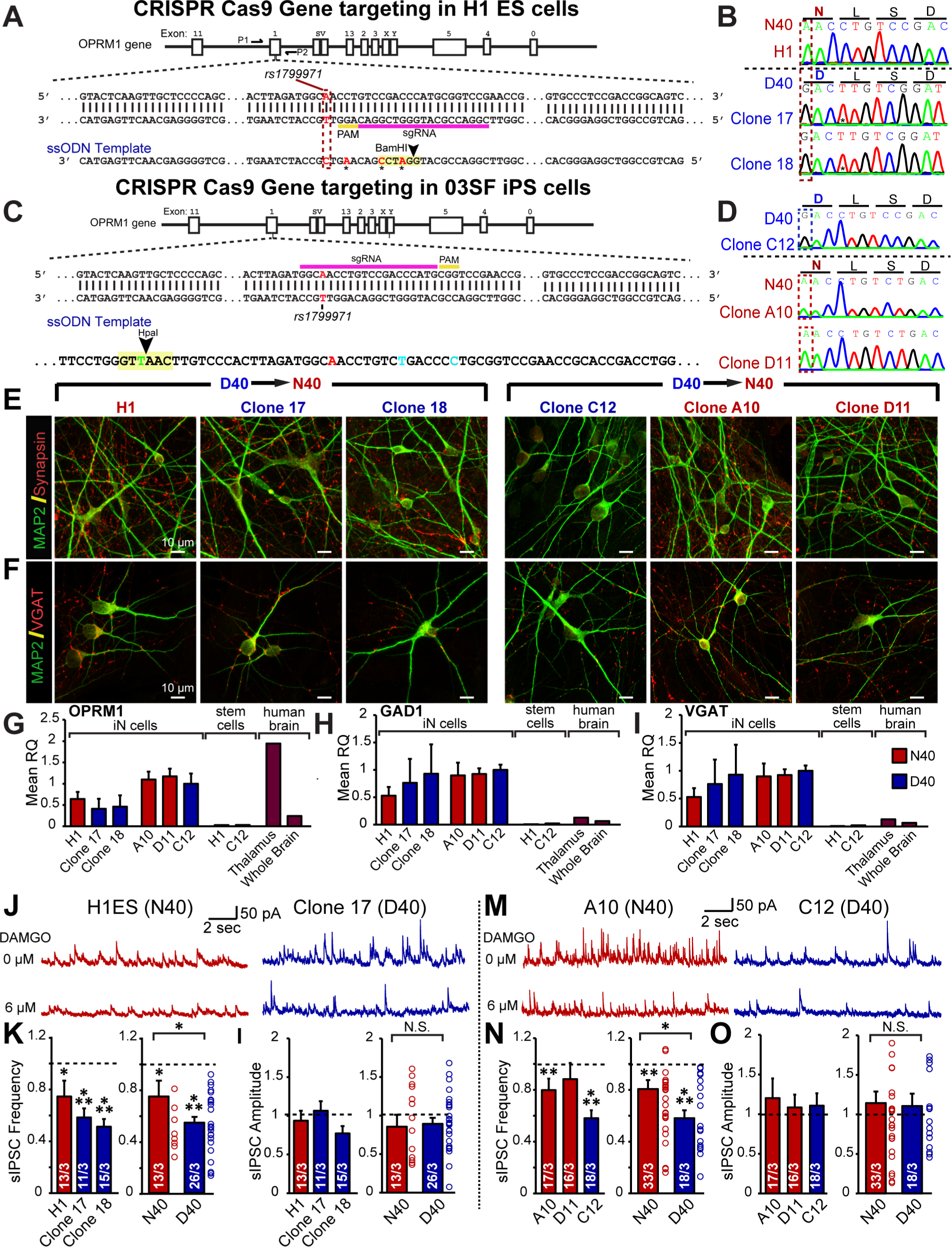
Human neurons from two sets of independently targeted isogenic human stem cell lines for OPRM1 A118G validate differential DAMGO response observed in patient cell lines. **(A)** OPRM1 Targeting Strategy 1: Structure of OPRM1 gene on chromosome 6 and schematic overview of CRISPR/Cas9 gene targeting strategy to knock-in homozygous G118 alleles into human H1 embryonic stem cell (H1ES) in which sgRNA targets donor strand. In the 140bp ssODN, we inserted a T to C mutation to incorporate OPRM1 GG118, synonymous G to A for PAM mutation, and synonymous G to C and G to A mutations to create a BamHI restriction enzyme site**. (B)** Sequencing of original H1ES control cell line carrying homozygous A118 (N40) alleles, and two isolated clones carrying homozygous OPRM1 G118 (D40) alleles (Clone 9-2-17, Clone 9-2-18). **(C)** OPRM1 Targeting Strategy 2: Structure of OPRM1 gene on chromosome 6 and an independent CRISPR/Cas9 gene targeting strategy to correct 03SF patient line (originally homozygous G118 expressing MOR D40) to homozygous A118 (N40). We designed a 200 nt template strand to knock-in homozygous A118 alleles, containing mutations to generate a HpaI restriction enzyme site for screening **(D)** Sequencing of passage-matched, uncorrected 03SF patient cell line carrying homozygous D40 alleles (C12) and two gene-corrected clones (Clone A10, D11) carrying homozygous OPRM1 A118 (N40) alleles after subcloning**. (E)** ICC of MAP2 (green) and Synapsin (red) of iN cells produced from gene-targeted ES cells and iPS cells. **(F)** Immunofluorescence of MAP2 (green) and VGAT (red) of iN cells produced from gene-targeted ES cells and iPS cells**. (G-I)** Relative mRNA levels of OPRM1 as well as markers inhibitory subtype specificity (GAD1, VGAT) measured by quantitative RT-PCR; mRNA levels are normalized to Synapsin I. Data are represented as means of three independently differentiated batches of iNs from each patient iPS cell line**. (J)** Representative traces of sIPSCs recorded to increasing concentrations of DAMGO in N40 and D40 iN isogenic iN cells derived from ES cells. **(K-L)** Quantification of sIPSC frequency (H1 vs control: p<0.05, Clone 17 vs control: p < 0.001, Clone 18 vs control: p <0.001, N40 vs control: p < 0.05, D40 vs control: p <0.001, N40 vs D40: p <0.05) and amplitude in response to 6µM DAMGO (H1 vs control: N.S., Clone 17 vs control: N.S., Clone 18 vs control: N.S., N40 vs control: N.S., D40 vs. control: N.S., N40 vs D40: N.S.) **(M)** Representative traces of sIPSCs recorded to increasing concentrations of DAMGO in N40 and D40 iN isogenic iN cells derived from iPS cells. **(N-O)** Quantification of sIPSC frequency (A10: DAMGO vs control: p<0.01, D11: DAMGO vs control: N.S., C12: DAMGO vs control: p <0.001, N40: DAMGO vs control: p<0.01, D40: DAMGO vs control: p<0.001, N40 vs D40: p <0.05) and amplitude in response to 6µM DAMGO (A10: DAMGO vs control: N.S., D11: DAMGO vs control: N.S., C12: DAMGO vs control: N.S., N40: DAMGO vs control: N.S., D40: DAMGO vs. control: N.S., N40 vs D40: N.S.). Data are depicted as means ± SEM. Numbers of cells/Number of independently generated cultures analyzed are depicted in bars. Statistical significance was evaluated by Student’s t test (* p<0.05, **p<0.01, *** p<0.001).

**Figure 3.**
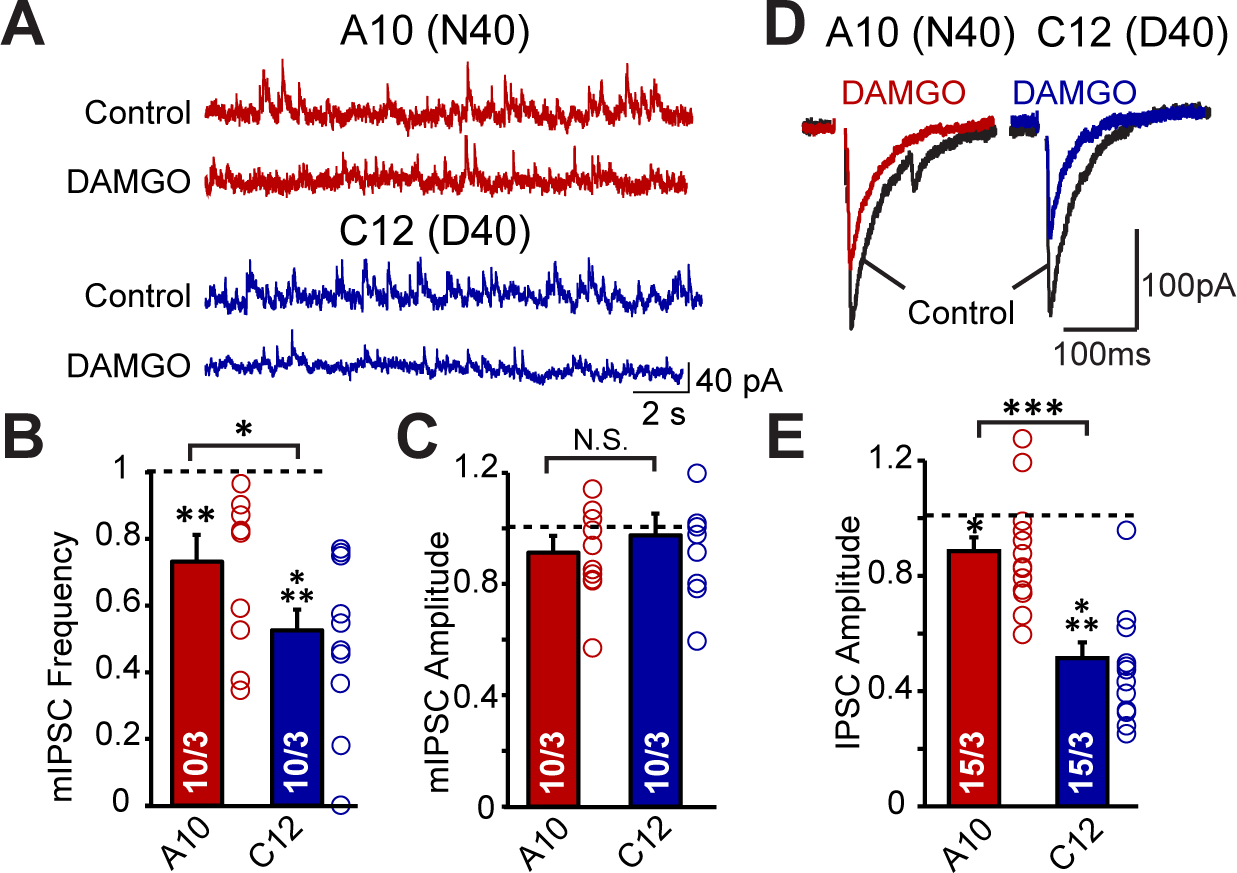
D40 iN cells exhibit greater inhibition of synaptic release and intrinsic excitability. **(a)** Representative traces of mIPSCs in one N40 (A10) and one D40 (C12) cell line derived iN cells recorded at 0mV holding potential and their response to DAMGO. **(b-c)** Quantification of mIPSC frequency (A10: DAMGO vs control p < 0.01, C12: DAMGO vs control p < 0.001) and amplitude (A10 and C12: DAMGO vs control p > 0.05) in A10 and C12 iN cells normalized to before DAMGO application. (DAMGO effect on frequency: A10 vs C12: p < 0.05, DAMGO effect on amplitude: A10 vs C12: N.S.) **(d)** Representative traces of evoked IPSCs from one N40 (A10) and one D40 (C12) cell line derived iN cells **(e)** Quantification of evoked IPSC amplitude in A10 and C12 iN cells normalized to before DAMGO application (A10: DAMGO vs control p < 0.05, C12: DAMGO vs control p < 0.001, A10 vs C12: p<0.001). Data are depicted as means ± SEM. Numbers of cells/Number of independently generated cultures analyzed are depicted in bars. Statistical significance was evaluated by Student’s t test (*p<0.05, **p<0.01, ***p<0.001).

### Isogenic human neurons recapitulate differential DAMGO response phenotype and exhibit altered synaptic function

In this highly controlled system of isogenic iN cells, we observed less culture-to-culture variability than the subject cell lines for OPRM1 mRNA and inhibitory neuronal markers (**Fig. 2G-I)**. Furthermore, we observed a similar decrease in sIPSC frequency compared to subject iN cells following acute DAMGO application, with a stronger inhibition in D40 versus N40 iN cells, and no effect on amplitude (**Fig 2J-O**). Furthermore, to determine whether the effect of DAMGO was mediated by MOR, we applied Naltrexone, a broad spectrum MOR antagonist, and found that the DAMGO-induced synaptic suppression could be reversed (**Supplemental Fig. 2J-K**). Thus, the reproducibility of the DAMGO response phenotype illustrates that the D40 variant alone explains the differential signaling and it is not due to secondary genomic variation.

We focused the remaining analyses on one pair of isogenic cell lines, C12 and A10, on the basis of their consistent differentiation and maturation. We found that DAMGO application more robustly decreases mIPSC frequency in D40 versus N40 iN cells, which no change in mIPSC amplitude (**Fig. 3A-C**), which suggests DAMGO mediated decrease in synaptic release. Consistent with the decrease in mIPSC frequency, we observed that DAMGO decreases evoked IPSC amplitude more robustly in D40 iN cells compared to N40 iNs (**Fig. 3D-E**). This is consistent with the hypothesis that MOR activation by DAMGO in human iN cells more robustly decreases neurotransmitter release probability in D40 iN cells compared to N40 iN cells, suggesting that the A118G SNP directly regulates synaptic function.

### D40 MOR-expressing neurons exhibit a more robust decrease in excitability following DAMGO compared to N40 iN cells

To understand whether the decreased synaptic release is compounded by decreased intrinsic excitability, we examined the effect of DAMGO on induced AP firing in N40 and D40 iN cells. We observed that 10 µM DAMGO induced D40 versus N40 iN cells to fire significantly fewer APs (**Fig. 4A-B**), with no effect on AP amplitude, firing threshold (**Fig. 4C-D**) or other properties including Time to reach peak or threshold (not shown). This is supported by an immediate and more robust decrease in spontaneous AP firing frequency following DAMGO application in D40 versus N40 iN cells (**Fig. 4E**), an effect which is sustained over the course of several minutes (**Fig. 4F**). This sustained decrease in AP frequency is paralleled by a rapid hyperpolarization of N40 and D40 iN cells (**Fig. 4G)**. This effect was found to be significantly more robust in D40 versus N40 iN cells in the first minute following DAMGO application. The immediate drop in AP firing frequency and membrane potentials in iN cells suggests that this may be occurring through a G-protein mediated signaling mechanism, which is activated immediately following agonist binding (21). No differences in AP rise time, decay time or half width were detected by DAMGO application (not shown). However, we observed a slight trending increase in the after hyperpolarization potential in the D40 versus N40 iN cells (**Fig. 4H-I**), with no significant difference in firing threshold or AP half width (**Fig. 4J-K**). These data indicate the functional differences between the two genotypes are at least partly mediated by a preferential decrease in excitability in D40 versus N40 iN cells, likely mediated by alterations in the G-protein coupled signaling cascade. Overall, these data suggest that DAMGO-induced decrease in excitability is superimposed by a synapse-specific effect, i.e., a stronger reduction in synaptic release probability mediated by presynaptic MOR at the nerve terminal in D40 MOR inhibitory neurons.

**Figure 4.**
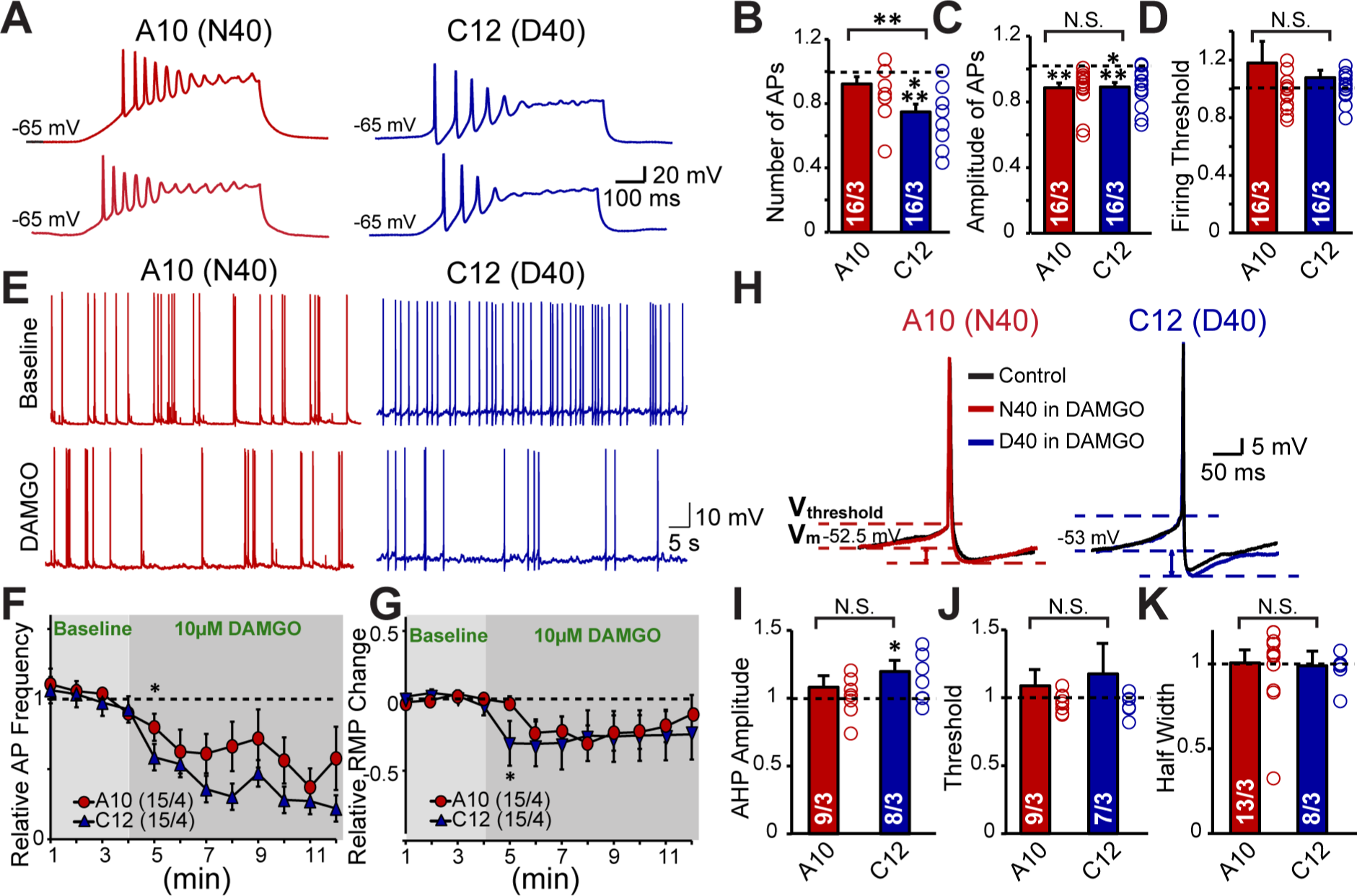
D40 iN cells exhibit a sustained decrease in intrinsic excitability. **(a)** Representative traces of repetitive action potentials generated from depolarizing current injections in one N40 (A10) cell line derived iN and one D40 (C12) cell line derived iN, and their response to DAMGO **(b-d)** Summary graphs of DAMGO effect on AP Number (A10: DAMGO vs control: N.S., C12: DAMGO vs control p<0.001), Amplitude (A10: DAMGO vs control p<0.01, C12: DAMGO vs control p<0.001) and firing threshold (A10: DAMGO vs control: N.S., C12: DAMGO vs control: N.S.). Data normalized to before DAMGO application reveals DAMGO preferentially decreases intrinsic excitability of D40 iNs but not N40 iNs (DAMGO effect on Frequency: A10 vs C12 p<0.01) with no effect on Amplitude (A10 vs C12: N.S.) or Firing Threshold (A10 vs C12: N.S.) **(e)** Representative traces depicting the effect of DAMGO on spontaneous action potential firing in one D40 (C12) and one N40 (A10) cell line derived iN **(f)** Quantification of number of spontaneous action potentials fired before and after DAMGO application in N40 and D40 iNs represented as a timecourse **(g)** Quantification of resting membrane potential before and after DAMGO application in N40 and D40 iNs represented as a timecourse **(h)** Representative traces of individual Action Potentials before and after DAMGO in N40 and D40 iN cells **(i-k)** Summary graphs depicting that DAMGO causes a trending increase in the AHP amplitude in D40 iN cells compared to N40 iN cells (A10: DAMGO vs control: N.S., C12: DAMGO vs control: p<0.05, A10 vs C12: N.S.) with no effect on Firing threshold (A10: DAMGO vs control: N.S., C12: DAMGO vs control: N.S., A10 vs C12: N.S.) or half width (A10: DAMGO vs control: N.S., C12: DAMGO vs control: N.S., A10 vs C12: N.S.). Data are depicted as means ± SEM. Numbers of cells/Number of independently generated cultures analyzed are depicted in bars. Statistical significance was evaluated by Student’s t test (*p<0.05, **p<0.01, ***p<0.001).

## DISCUSSION

Our study provides the first experimental evidence detailing the electrophysiological consequences of the N40D SNP on MOR activation in its endogenous human neuronal context. First, we generated iN cells from human subject-derived stem cells carrying homozygous alleles for N40 MOR or D40 MOR and found that D40 MOR expressing iN cells exhibit stronger inhibitory effects of MOR activation on synaptic release. Second, to validate the functional consequences of the SNP in a system highly controlled for background genetic variation, we used CRISPR/Cas9 mediated gene targeting to: 1) knock-in homozygous D40 alleles into H1ES cells; 2) correct the homozygous D40 alleles in 03SF iPS cell subject line into N40 alleles, and thus generated two sets of isogenic stem cell lines for highly controlled mechanistic analyses, The isogenic iN cells not only recapitulated the DAMGO response phenotype of the patient iN cells, but also revealed that the N40D SNP mediates a more robust decrease in excitability and synaptic release.

Despite previous studies in knock-in mouse models and heterologous expression systems, the precise molecular and cellular consequences of MOR N40D have remained unclear, primarily due to species-specific and context-specific mechanisms in the modulation of MOR signaling. For instance, rodent models and other expression systems have suggested that the D40 allele confers a “gain-of-function” effect by causing increased potency for DAMGO and other MOR agonists (7, 11, 12, 22). However, subsequent studies have reported that the D40 allele is associated with reduced mRNA and protein expression in multiple brain regions of knock-in mice (10, 13) along with reduced antinociceptive responses to morphine (8), providing support for a “loss-of-function” phenotype. These contradictory results strongly necessitate the need for a human neuronal model to understand MOR function.

The novelty of our approach using human iN cells to investigate the synaptic pathology of addiction is that these cells carry the genetic signatures of the subjects from whom they were derived. Identifying identical DAMGO-mediated responses across multiple subject-derived and CRISPR-edited cell lines generated using independently executed targeting strategies clearly demonstrates that the observed effect is a direct consequence of only the MOR N40D variant. Collectively, this overall approach of combining multiple patient lines with genome-engineered isogenic lines to assess addiction-associated electrophysiological phenotypes has not been fulfilled in previous work. Thus, we show the utility of disease modeling using stem cell derived disease-relevant cell types as a framework to the field of addiction for conducting future mechanistic analyses.

This study represents a significant advance in our understanding of the neurobiological mechanisms underlying the human N40D MOR variant in a human neuronal context. Our study provides direct evidence that common genetic variation encodes functional variation at the level of synaptic transmission. The use of patient-derived stem cells to unravel the impact of OPRM1 gene variants may ultimately provide the necessary insight to develop patient-specific, precision medical interventions for drug and alcohol dependence.

## ACKNOWLEDGMENTS

We thank RUCDR Infinite Biologics for generating the iPS cells from human subjects and assisting with CRISPR/Cas9 gene targeting on 03SF iPS cell line. Research is supported by grants from NIH-NIAAA R01 AA023797 as well as Collaborative Studies on the Genetics of Alcoholism/COGA 5U10AA008401-26. AH is supported by NIH-NIAAA NRSA F31AA024033. We are grateful to the members of the Collaborative Genetic Study of Nicotine Dependence (COGEND) for the selection of human subjects, and we are grateful to the de-identified individuals who contributed tissue to the study.

## Supplemental Information

**Supplementary Figure 1.**
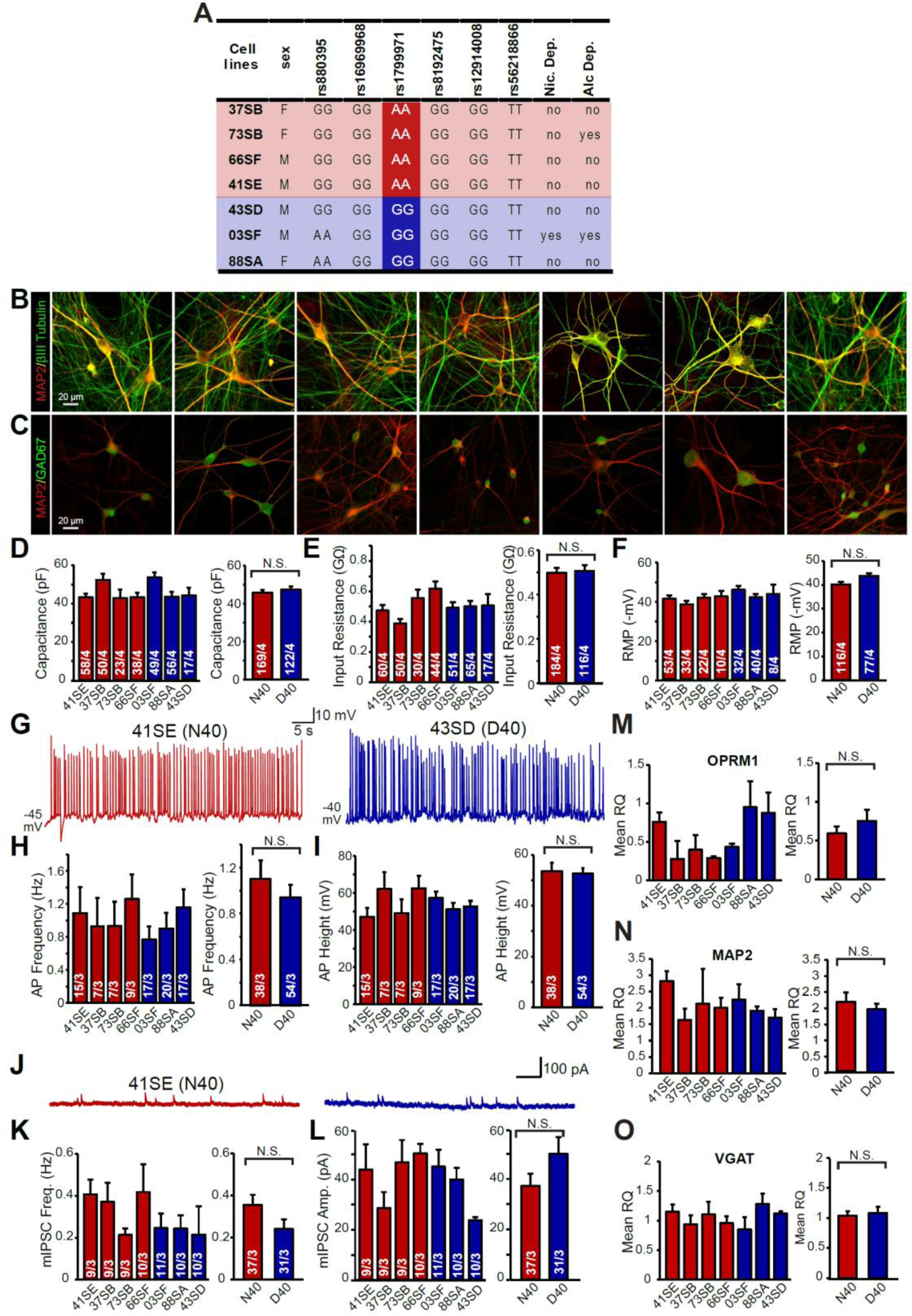
OPRM1 N40D SNP does not impair intrinsic neuronal parameters of iN cells, inhibitory neuronal differentiation, or neuronal function. **(A)** iPS cell lines generated by RUCDR Infinite Biologics from individuals carrying A118G SNPs are homozygous for SNPs in other genes linked to addiction. **(B)** MAP2 and βIII-tubulin ICC of inhibitory iN cells differentiated from human iPS cell subject lines to illustrate expression of markers of neuronal maturation **(C)** MAP2 and GAD67 immunofluorescence of inhibitory iN cells differentiated from human iPS cell subject lines to illustrate inhibitory subtype **(D-F)** N40 and D40 iPS cell derived iN cells exhibit similar intrinsic membrane properties including capacitance (N40 vs D40: N.S.), input resistance (N40 vs D40: N.S.), and resting membrane potential (N40 vs D40: N.S.), illustrating similar maturation status **(G-I)** Representative traces of spontaneous action potentials recorded from one A118 (N40) and one G118 (D40) iN cells. Summary graphs illustrate that N40 and D40 iNs exhibit similar spontaneous action potential firing frequency (N40 vs D40: N.S.) and amplitude (N40 vs D40: N.S.) **(J-L**) Frequency (N40 vs D40: N.S.) and amplitude (N40 vs D40: N.S.) of miniature IPSCs in iN cells are unaffected by MOR N40D substitution **(M-O)** Relative mRNA levels of OPRM1 (N40 vs D40: N.S.) as well as markers of neuronal maturation including MAP2 (N40 vs D40: N.S.) and Tuj1 (N40 vs D40: N.S.) measured by quantitative RT-PCR; mRNA levels are normalized to Synapsin I. Data are represented as means of three independently differentiated batches of iNs from each patient iPSC line. Summary graphs of these parameters are shown as merged data of either four N40 patients or three D40 patients. Data are depicted as means ± SEM. Numbers of cells/Number of independently generated cultures analyzed are depicted in bars. Statistical significance was evaluated by Student’s t-test (* p<0.05, ** p<0.01, *** p<0.001)

**Supplementary Figure 2.**
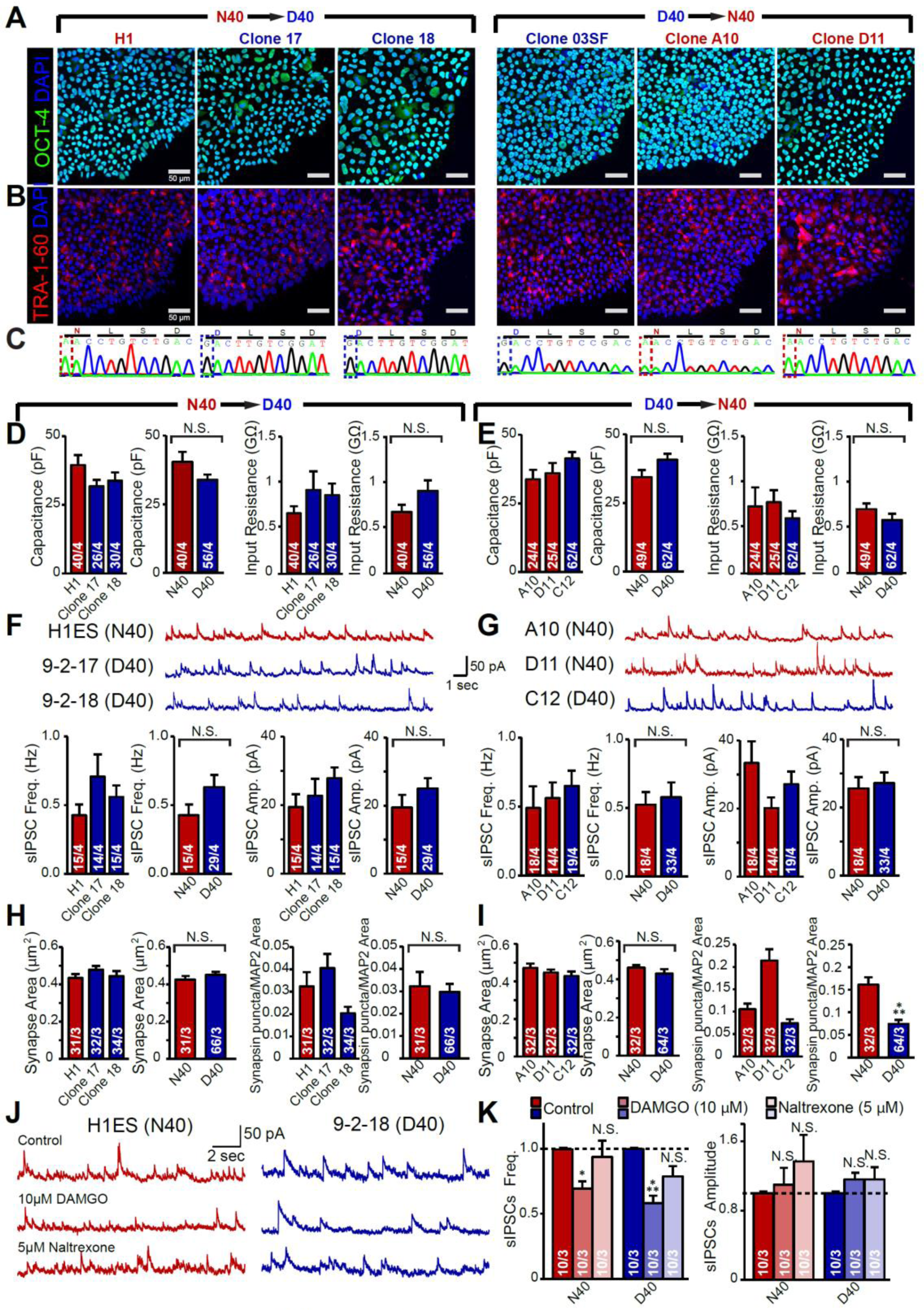
Characterization of CRISPR/Cas9 gene targeted isogenic cell lines. **(A)** Oct4 (green) and Dapi (blue) immunofluorescence for H1ES and 03SF gene-targeted clones depicting pluripotency **(B)** Tra-1-60 (red) and Dapi (blue) immunofluorescence for H1ES and 03SF gene-targeted clones depicting pluripotency **(C)** Sequencing of original H1ES control and two homozygous MOR D40 lines confirming homozygous knock-in of either N40 alleles or D40 alleles. **(D)** N40 and D40 isogenic iN cells from H1 ES cells exhibit similar intrinsic membrane properties including capacitance (N40 vs D40: N.S.) and input resistance (N40 vs D40: N.S.) at baseline levels, illustrating similar maturation status **(E)** N40 and D40 isogenic iN cells from 03SF iPS cell line exhibit similar intrinsic membrane properties including capacitance (N40 vs D40: N.S.) and input resistance (N40 vs D40: N.S.) at baseline levels, illustrating similar maturation status **(F-G)** Both N40 and D40 iN cells exhibit sIPSCs; Frequency (H1 ES cell isogenic lines: N40 vs D40: N.S., 03SF iPS cell isogenic lines: N40 vs D40: N.S.) and amplitude (H1 ES cell isogenic lines: N40 vs D40: N.S., 03SF iPS cell isogenic lines: N40 vs D40: N.S.) of sIPSCs in iN cells are unaffected by MOR N40D substitution **(H-I)** Synapse area (H1 ES cell isogenic lines: N40 vs D40: N.S., 03SF iPS cell isogenic lines: N40 vs D40: N.S.) and Synapsin puncta density (H1 ES cell isogenic lines: N40 vs D40: N.S., 03SF iPS cell isogenic lines: N40 vs D40 p<0.001) normalized to MAP2 area are unaffected by MOR N40D substitution **(J-K)** Naltrexone reverses the DAMGO synaptic inhibition phenotype in both N40 and D40 iN cells (N40 Effect on Frequency: DAMGO vs control p < 0.05, Naltrexone vs control: N.S., D40 Effect on Frequency: DAMGO vs control p <0.001, Naltrexone vs control: N.S.) with no effect on amplitude (N40 Effect on Amplitude: DAMGO vs control: N.S., Naltrexone vs control: N.S., D40 Effect on Amplitude: DAMGO vs control: N.S., Naltrexone vs control: N.S.). Data are presented as mean ± SEM, Numbers of cells/Number of independently generated cultures analyzed are depicted in bars. Student’s t-tests were used for statistics.

**Supplementary Figure 3.**
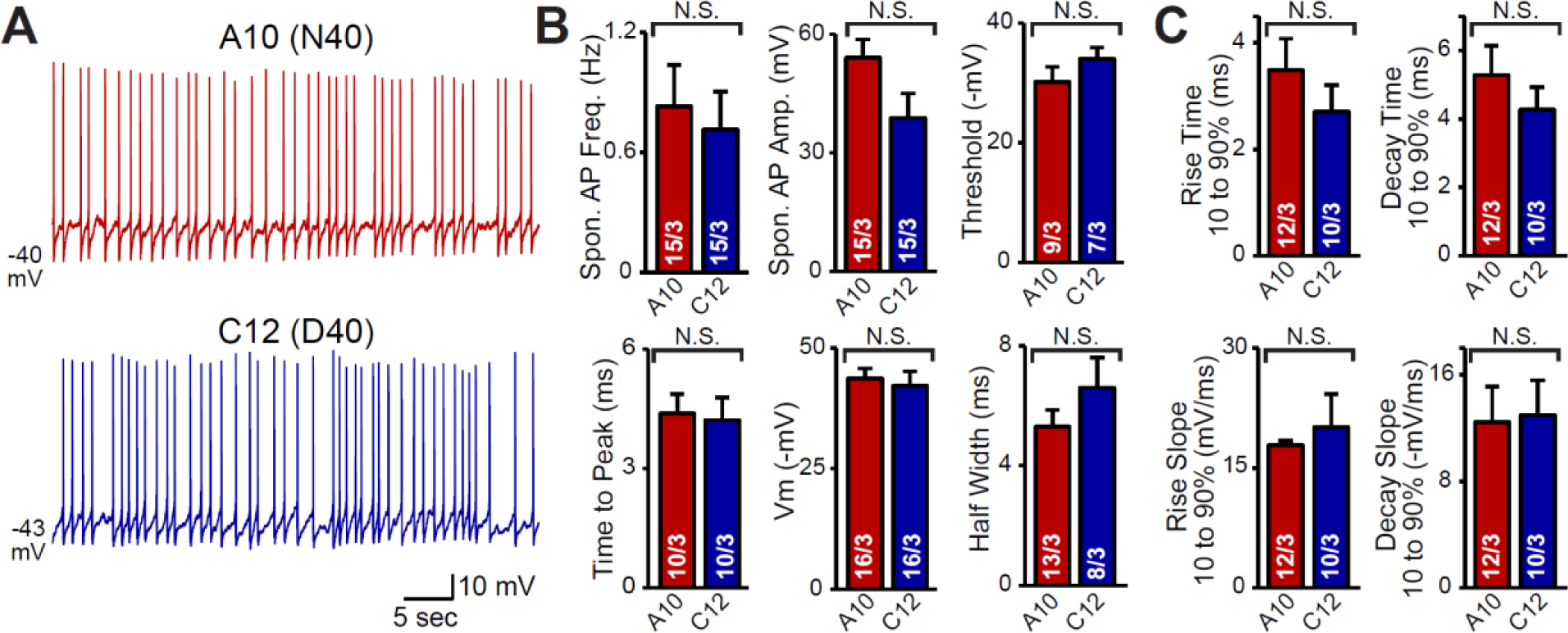
MOR N40D SNP does not impair action potential properties in N40 and D40 iN cells at baseline levels. **(A)** Representative traces of spontaneous action potentials fired by A10 (N40) iN cells and C12 (D40) iN cells **(B)** Quantification of action potential properties at baseline levels between A10 (N40) and C12 (D40) iN cells shows N40D does not alter Spontaneous AP frequency (A10 vs C12: N.S.), amplitude (A10 vs C12: N.S.), threshold (A10 vs C12: N.S.), time to peak (A10 vs C12: N.S.), resting membrane potential (A10 vs C12: N.S.), and half width (A10 vs C12: N.S.) **(C)** Quantification of action potential rise and decay kinetics at baseline levels between A10 (N40) and C12 (D40) iN cells shows N40D does not alter Rise Time (A10 vs C12: N.S.), Decay time (A10 vs C12: N.S.), Rise slope (A10 vs C12: N.S.), or decay slope (A10 vs C12: N.S.). Data are depicted as means ± SEM. Numbers of cells/Number of independently generated cultures analyzed are depicted in bars. Statistical significance between N40 and D40 was evaluated by Student’s t-test (*p<0.05, **p<0.01, ***p<0.001)

